# Emotion arousal but not valence is strongly represented in aperiodic EEG activity stemming from thalamocortical interactions

**DOI:** 10.1101/2024.03.11.584477

**Authors:** Ritu M Borah, Anagh Pathak, Arpan Banerjee

## Abstract

Oscillatory EEG signals contain both periodic and aperiodic (1/f components), segregating which is important for gaining insights into the underlying neural computations. Several studies have explored the periodic features underlying neural basis of emotion processing, while 1/f component is less explored. Hence, the present study sought to determine whether seeing emotionally laden music video clips have an impact on aperiodic components along with the periodic components and their underlying basis by conceptualizing a biophysically realistic computational model. Using the publicly available Data for Emotional Analysis using Physiological Signals (DEAP) dataset we elucidate that there is a significant increase in exponent and offset of aperiodic background during high arousal compared to low arousal trials. In contrast, there were no discernible variations between the aperiodic components during the high and low valence categories. In high arousal, we also discovered reduction in alpha power in line with previous studies. Finally, to elucidate the underlying mechanisms, we employed a biophysically realistic and previously established thalamo-cortical model of neural field activity, which which allowed us to characterize that heightened emotional arousal appears emerge from an increased cooperation in information communication between neuronal populations in reticular and relay nuclei of thalamus.

**Significance statement:** Dissociation of emotion and valence and arousal is an established theory in affective neuroscience. Are these dimensions represented in aperiodic background of macrocscopic electrophysiological scales of EEG? We sought to ask this question in the publicly available DEAP data. Our findings of increased aperiodic components during high arousal in contrast to low arousal suggest heightened neural activity alongside an inhibitory influence. Using a neural field model of thalamo-cortical interactions reveals a mechanistic approach that suggests heightened arousal corresponds to an enhanced inhibitory coupling between thalamic reticular and relay neuronal populations. In essence, this study advances our understanding of the intricate relationship between electrophysiological signals and emotion dimensions, shedding light on the mechanisms underlying arousal-related emotional responses.

## Introduction

Emotions play a pivotal role in modulating human perception and action (Zadra & Clore, 2011), attention (Carretié et al., 2001; Dolcos et al., 2020), enhancing memory (Tyng et al., 2017) (Dunsmoor et al., 2019)), social and economic decision-making (Eijlers et al., 2019). Hence, understanding the neurophysiological mechanisms underlying our emotional experiences offers a unique perspective to investigate various cognitive functions. A well-established conceptual framework, identifies two dimensions - valence and arousal, along which emotional states can be quantified (Posner et al., 2005; Russell, 1980)). Valence pertains to the spectrum of unpleasant to pleasant emotional experiences, while arousal relates to the degree of excitement associated with an emotional state. Several studies have employed electroencephalogram (EEG) paradigms involving stimuli such as facial expressions, music, and virtual reality experiences to induce emotions and explore their neural correlates (Costa et al., 2006; Daly et al., 2014; De Cesarei & Codispoti, 2011; Hofmann et al., 2021). For instance, a decrease in alpha power has been consistently associated with high arousal states (Hofmann et al., 2021; Kim et al., 2021; Schubring & Schupp, 2021). However, it is worth noting that a contrary finding of increase in alpha power correlated with increased arousal also exist (Aftanas et al., 2004; Uusberg et al., 2013). Alpha power decrease is also observed with negative emotions e.g., fear (Eijlers et al., 2019). Additionally, beta power reductions have been linked to increased arousal (Kim et al., 2021; Schubring & Schupp, 2021), while frontal midline theta (4-7Hz) activity has been correlated with pleasant stimuli (Sammler et al., 2007). While, these studies highlight the significance of rhythmic oscillations in understanding the neural underpinnings of emotion, an explanation of the disparate findings related to alpha increase or decrease remains elusive.

A major thrust in neural signal processing has currently shifted to the understanding of both rhthymic and arrythmic (non-oscillatory or 1/f) components (He, 2014; He et al., 2010) the latter being hypothesised to confound the measurement of true neural oscillations (Gyurkovics et al., 2021; Voytek et al., 2015). Intriguingly, researchers have also proposed that the 1/f activity is a hallmark of criticality (Buzsáki, 2006) - the neurobiological state of homeostasis among excitatory and inhibitory populations (Donoghue et al., 2020; Gao et al., 2017; Voytek et al., 2015) maintained in a putative brain region to process incoming stimuli. Supporting this possibility, emerging evidence suggests that, arrhythmic components also hold relevance as correlates of cognitive processes, e.g., predicting cognitive processing speed (Ostlund et al., 2021; Ouyang et al., 2020), charting the lifespan trajectories of cognitive processing (Thuwal et al., 2021), thus necessitating a comprehensive approach to their analysis (Colombo et al., 2019; Lendner et al., 2020; Voytek et al., 2015). Finally, the accurate characterization of the arrythmic activity can also reconcile the disparate observations, such as alpha decrease/ increase in different cohorts(Gyurkovics et al., 2021). Consequently, it is imperative to investigate the role of aperiodic brain dynamics in shaping emotional states—an area that has thus far remained scantly explored in the field of emotion research.

From a rigorous neurbiological perspective, numerous studies have illuminated the intricate involvement of subcortical regions—the thalamus, amygdala, and various sensory and frontal cortical regions—in processes related to emotional arousal and valence processing (Karjalainen et al., 2019). Notably, fMRI investigations consistently reveal a positive correlation between amygdala activation and emotionally arousing stimuli, irrespective of their valence (Lin et al., 2020) as well as existence of multitasking core systems that process emotion (Barrett, 2017) (Barrett, 1998)(Barrett, 2017). Interestingly, excitatory connection from the amygdala to thalamic reticular nuclei, serving as an attentional regulatory mechanism, was identified in primates (Barbas et al., 2011). These insights have led us to hypothesize that the neural mass model involving dynamic interactions between thalamus and cortex ((Robinson et al., 2003) (Robinson et al., 1997), which has been frequently applied in various brain function contexts (e.g., arousal, epileptic seizures, sleep-wake cycle (Rennie et al., 2002), may offer valuable insights into the complex aperiodic dynamics of emotional arousal. Most importantly, this model can generate both aperiodic and periodic spectra of EEG, making it a potent instrument for simulating and comprehending underlying brain activity. We specifically posit that increased emotional arousal modulates the thalamic reticular nuclei, thereby facilitating enhanced/ decreased inhibitory coupling with the relay nuclei which in turn feeds back to cortical population and govern EEG 1/f spectra. Thus, our main objectives are two-fold: first, to characterize EEG from the DEAP (Data for Emotional Analysis using Physiological Signals) database (Koelstra et al., 2012) in aperiodic and periodic components across the dimensions of valence and arousal, and second, to explore the extent to which the thalamocortical neural mass model can account for these observations.

## 2. Methods

### 2.1 Dataset and Experimental Design

We utilized the Open Source affective Database for emotion analysis using Physiological signals (DEAP) (Koelstra et al., 2012). The dataset includes EEG recordings (at a sampling rate of 512Hz, using Biosemi Active Two system) and physiological signals such as blood volume pressure, respiration rate, and temperature. The data was collected from 32 participants, with an equal distribution of male (16) and female (16) participants. The participants’ mean age was 26.9 years, ranging from 19 to 37 years.

During the experiment, each participant watched forty different 60-second music videos. The presentation of the videos was separated by a fixed inter-trial interval of 5 seconds. After viewing each video, there was a 3-second fixation window, and then the participants were asked to rate their emotional experiences using continuous 1-9 Likert scales for valence, arousal, dominance, and liking. Additionally, participants rated their familiarity with the videos on a Likert scale, 1-4 . For the mentioned rating, relevant self-assessment manikins were displayed in the middle of the screen. The participants rated their experience by moving the mouse below the manikins in a continuous scale. They were instructed to rate their valence from feeling of unhappy or sad to happy or joyful state. The arousal state from calm or bored to stimulated or excited. For the ratings, they were instructed to define their interests of like-dislike and not of feelings in particular. Since we were interested in particular in valence and arousal states, we only extracted only those two behavioural rating for our study.

### 2.2 Preprocessing

We used EEGLAB for the preprocessing of the data. Initially, a bandpass filter with a range of 2-45 Hz (pop_eegfiltnew.m) was applied. Subsequently, bad channels were removed using cleanline function where a channel was removed if there is 5s flatline, standard deviation of high frequency signals is greater than 5 and the correlation to the nearest channel is less than 0.8. Epochs were extracted, spanning from -6 to 61 seconds from the onset of video stimuli and then Independent Component Analysis (ICA) was performed (pop_runica). Artifacts were labeled using default ICA label algorithm and a component was rejected if it was identified as eye, muscle, heart, channel with 90% accuracy. To establish a common reference, the data was re-referenced to the average (pop_reref).

The Multitaper method (mtspectrumc, CHRONUX) was employed to convert the time series data into the frequency domain. Prior to conducting the multi-taper analysis, detrending was was done (detrend.m). The power spectrum for each trial and channel was then estimated.

To conduct the multi-taper analysis, a taper bandwidth of 0.6 Hz [36 (60s×0.6s) time bandwidth product and 45 tapers]. Additionally, the tapers bandwidth was kept consistent with the 5-second pre-stimulus fixation period, which utilized 5 tapers.

### 2.3 Estimation of Aperiodic and Periodic components

Fitting Oscillations and One Over F (FOOOF) toolbox in MATLAB was employed to parametrize the power spectra of the EEG data into periodic and aperiodic components (Donoghue et al., 2020)))). The FOOOF algorithm involves several steps to fit and analyze the spectra. Initially, it fits an aperiodic component, which is represented as a straight line when plotted in log-log frequency and power space, unless a knee or break in the spectrum is predicted or specified. This fitted aperiodic component is then subtracted from the original raw spectra, resulting in residuals. Next, the periodic and noise components are estimated based on these residuals. To identify peaks in the spectrum, a Gaussian function is fitted to the residual spectrum, and a peak is detected when the fitted function surpasses the noise threshold. The process of fitting the Gaussian is performed iteratively, depending on either reaching the maximum peak detection limit set or when the next Gaussian falls below the noise threshold. The final raw spectrum is obtained by fitting the Gaussian function(s) along with the aperiodic component, yielding a comprehensive representation of the entire spectrum. This allows for the separation and extraction of rhythmic and arrhythmic components.

For modelling, the power spectrum (*PSD*) was defined as the mix of Gaussian and Lorentzian function as in equation (1)

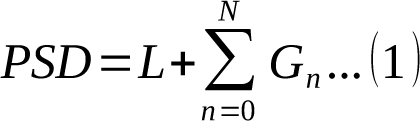

The Gaussian (*G_n_*) is defined as function of peak power (*a*) in log_10_ (*power*), central frequency (*c*), standard deviation (*ω*) and vector of input frequencies (*F*) as follows

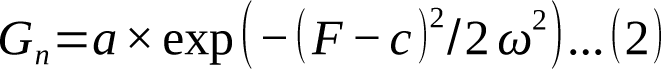

The Lorentzian (*L*) giving the aperiodic fit is explained by the broadband offset (*b*), exponent (*Χ*) and knee parameter (*κ*) which determine if the bend or break in the slope of the power spectrum

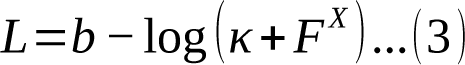

For the model fit, following parameters were applied. The frequency range was set to [4 40Hz], the maximum number of peaks allowed was 5 for the 60-second stimulus fit and 5 for the 5-second pre-stimulus fit. The minimum peak amplitude was required to exceed the aperiodic fit, set at 0.0. The aperiodic mode was fixed, and the peak width limits were restricted to [0.5 12 Hz]. Additionally, a peak threshold of 2.0 was used.

This procedure was employed on the power spectrum of average of electrode for each trial. The estimated exponent, offset, and peak alpha power were used for further analysis. In instances where multiple alpha peaks were detected, the one with the maximum power was selected for subsequent analysis. We subsequently fitted the model on each channel level to investigate the topography of the spectral parameters (Fig 2).

**Figure 1:**
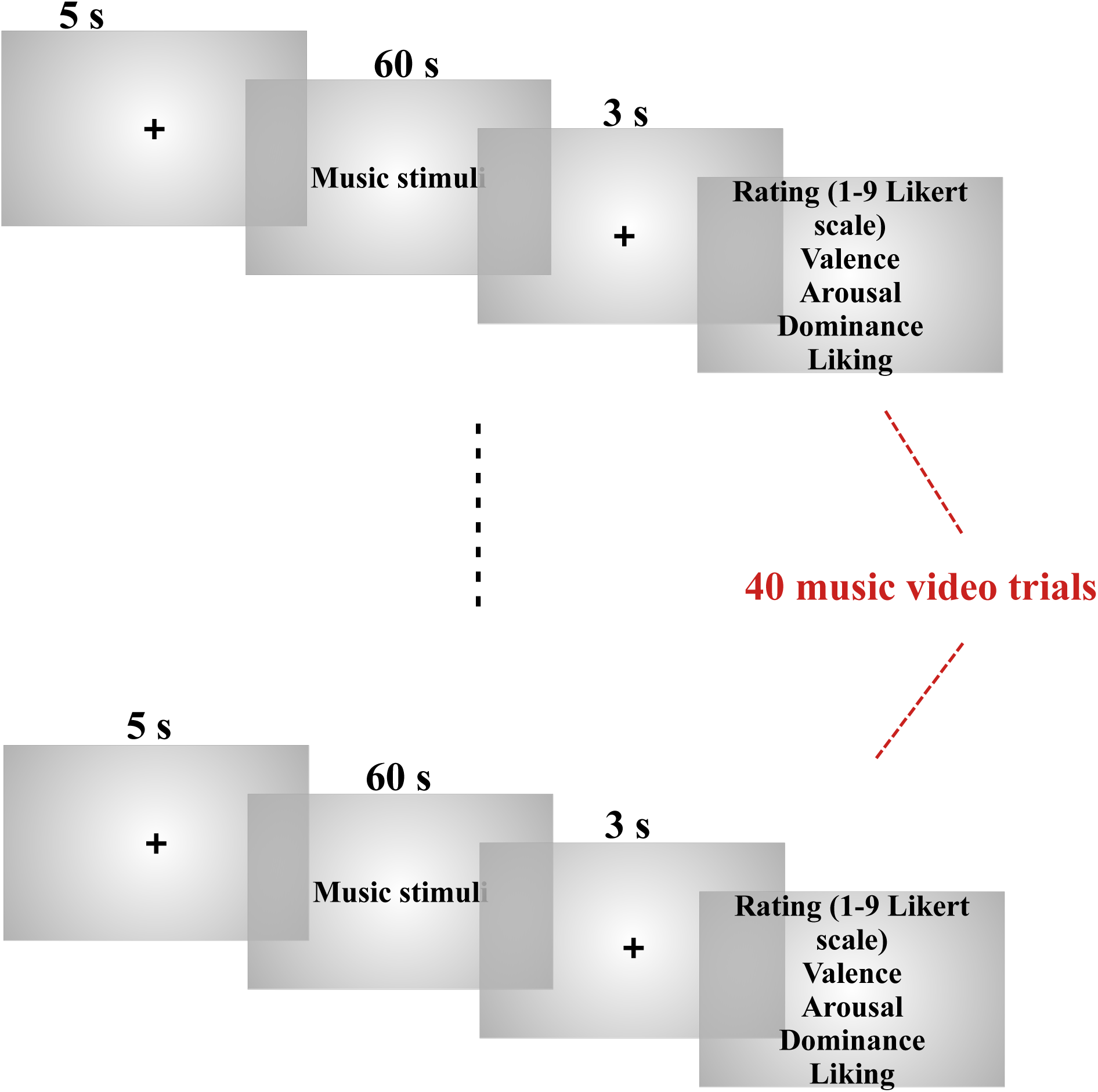
Pictorial representation of stimulus presentation: one trial involved a pre stimulus fixation of 5s, followed by 60s music video (randomly presented to participants) and 3s fixation window. After which participant gives their rating for Arousal, valence (also dominance, liking and familiarity; not considered for our current study).

**Figure 2:**
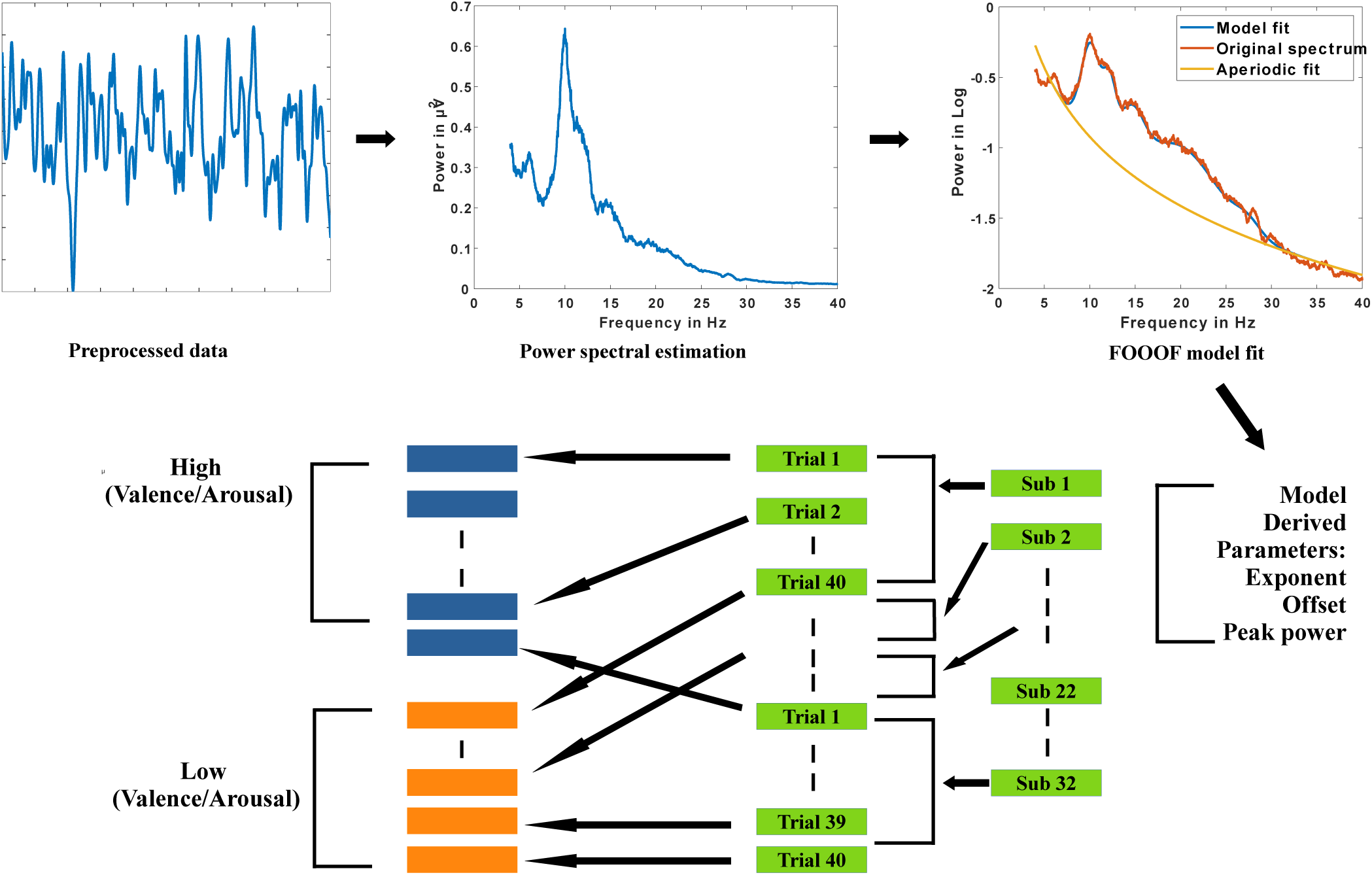
Illustration of the FOOOF application ((Donoghue et al., 2020)) to obtain high and low states of valence and arousal separately from the DEAP dataset.

### 2.4 Statistical paradigm

For each video, participants were asked to rate in 1-9 Likert scale their arousal and valence states. Each video was considered as a trial and based on the ratings given were divided into high-low arousal and valence categories. The threshold for high—low division was kept at 5 adapted from the original work (Koelstra et al., 2012)).

We employed permutation method to evaluate the difference in means between the high—low (exponent, offset, alpha power) arousal and valence conditions because we wanted to bypass the Gaussian assumption of the data. We made 20 runs on random 500 trials of each condition with 5000 iterations on each run. The function follows the principle of randomly shuffling data and calculating t-score, followed by p-value estimation. In addition, the effect size was computed as the observed difference divided by the mean of the standard deviations of the two samples.

We used MATLAB function randanova2 taking valence (High/Low) and arousal (High/Low) as factors. The function takes in the grouped data and calculates F values from the original data and the randomly shuffled data for the provided iterations (5000). The p-value was estimated as proportion of F-value of randomly shuffled data that are equal to or greater than the original F-value. The method uses ANOVAN model fit without the assumption of Gaussian data distribution making it more robust. We also tested for the interaction effect between arousal and valence.

The complete analysis was done in subject independent manner i.e. all trials of subjects were clubbed based on condition, to increase statistical power because there were only few trials per participant on performing the binary classification of high and low states.

### 2.5 Neural mass model of EEG dynamics

In order to arrive at a mechanistic explanation for observed spectral changes in ongoing recordings as a function of valence and arousal, we employed a well-established neural mass modelling approach of EEG/MEG dynamics that incorporates the thalamocortical interactions. The model consists of a recurrent population of cortical and thalamic neurons where the average dynamics of each population (neural mass) is described coupled rate equations (Fig 3)(Rennie et al., 2002; Robinson et al., 1997, 2003). The thalamus consists of reticular and relay populations. The excitatory relay cells convey ascending inputs to the cortex and are kept under inhibitory control by the reticular population (Fig 3) which receives inputs from structures like amygdala (Barbas et al., 2011) . The cortex is represented as an excitatory-inhibitory mass that receives subcortical input from the relay cells and feeds back to the thalamic populations. All thalamocortical interactions are subject to a finite transmission delay which is kept as 40 ms throughout the article, keeping in line with previous studies. In the absence of overt stimuli, the relay populations are fed by noisy input . Each population in the model is characterized by its mean soma membrane potential (V), mean firing rate (Q) and the local presynaptic activity (*ϕ*). Differential equations are employed to specify the conversion of these quantities, mimicking the effects of neuronal gain and synapto-dendritic filtering. The following second order differential equation specifies how the mean membrane potential (V) of a population changes in response to input presynaptic activity (*ϕ*)-

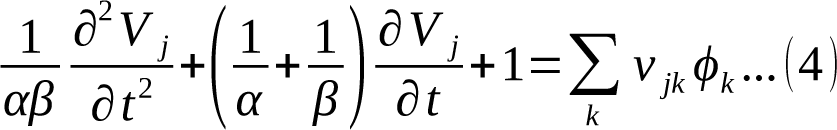

**Figure 3:**
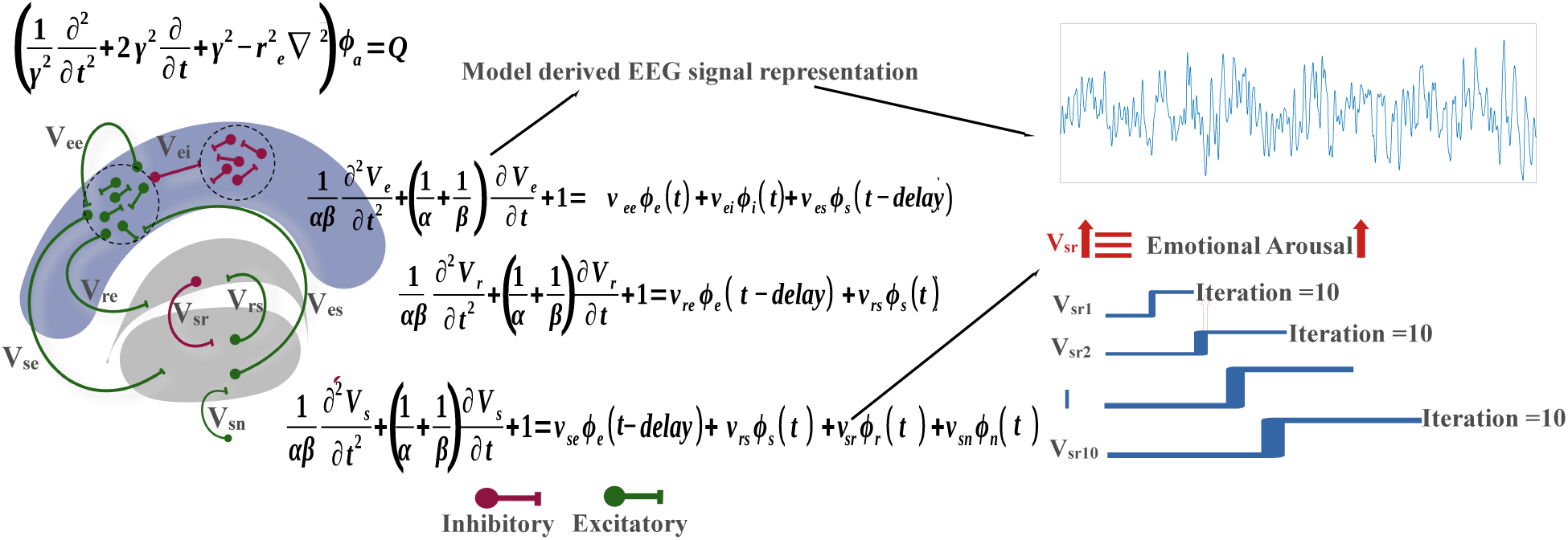
Schematic representation of the thalamocortical model showcasing feedforward and feedback loop between thalamus and cortex. Here, e is excitatory, i is inhibitory, s is relay nucleus of thalamus, r is reticular nucleus of thalamus, V_ab_ is membrane potential and the subscript denote the respective interaction between the neuronal population where ab can be e, i, s, r; ϕ_a_ is presynaptic potential

Where *α* and*β*are the inverse rise and decay times respectively of membrane potential generated by an input impulse. The resulting membrane potential (V) maps to the mean firing rate (Q) according to the following sigmoidal activation function-

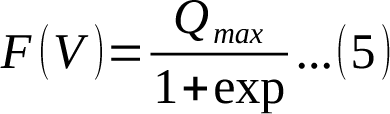

Where *Q_max_*is the maximum firing rate, *θ* is the mean firing threshold and *σ* is its standard deviation. Finally, the mean soma firing rate (Q) is converted to a resulting local presynaptic activity (*ϕ*) through the following damped wave equation-

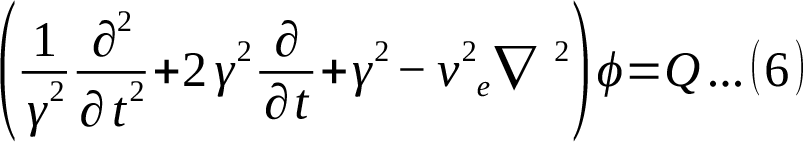

For normative parameter values (see Table 1), the model mimics the dynamics of resting state human EEG/MEG, with prominent peak in alpha (8-12Hz) frequency band, and a distinct 1/f broadband slope.

**Table 1:** Baseline resting state parameter values (adapted from Robinson et al., 2003; Robinson et al., 1997)

| Parameter | Resting state value |
| --- | --- |
| $\theta$ | 15mV |
| $\sigma$ | 3.3 mV |
| Q | $250s^{-1}$ |
| $\gamma$ | $100s^{-1}$ |
| $\alpha$ | $50s^{-1}$ |
| $\beta$ | $200s^{-1}$ |
| $t_0$ | 80 ms |
| $v_{ee}, v_{es}, v_{se}$ | 1.2 mVs |
| $v_{ei}$ | -1.8 mVs |
| $v_{sr}$ | -0.8mVs |
| $v_{sn}\phi_n$ | 1.0mVs |
| $v_{\mathfrak{R}}$ | 0.4mVs |
| $v_{rs}$ | 0.2mVs |

## Results

### Increase in exponent and offset, and decrease in peak alpha power in high arousal compared to low arousal

We assessed the impact of arousal and valence on exponent and offset of 1/f slope of EEG activity. To determine the statistical significance of these effects, we conducted permutation test for arousal and valence conditions separately. Here, we present these findings based on 20 runs as the mean ± standard deviation of the runs. For reporting the p-value and effect size, the maximum and minimum observations, respectively, are utilized. Additionally, to examine interaction between the two factors (arousal and valence) we performed ANOVA-based permutation tests for the spectral parameters (Fig 4, Table 2).

**Figure 4:**
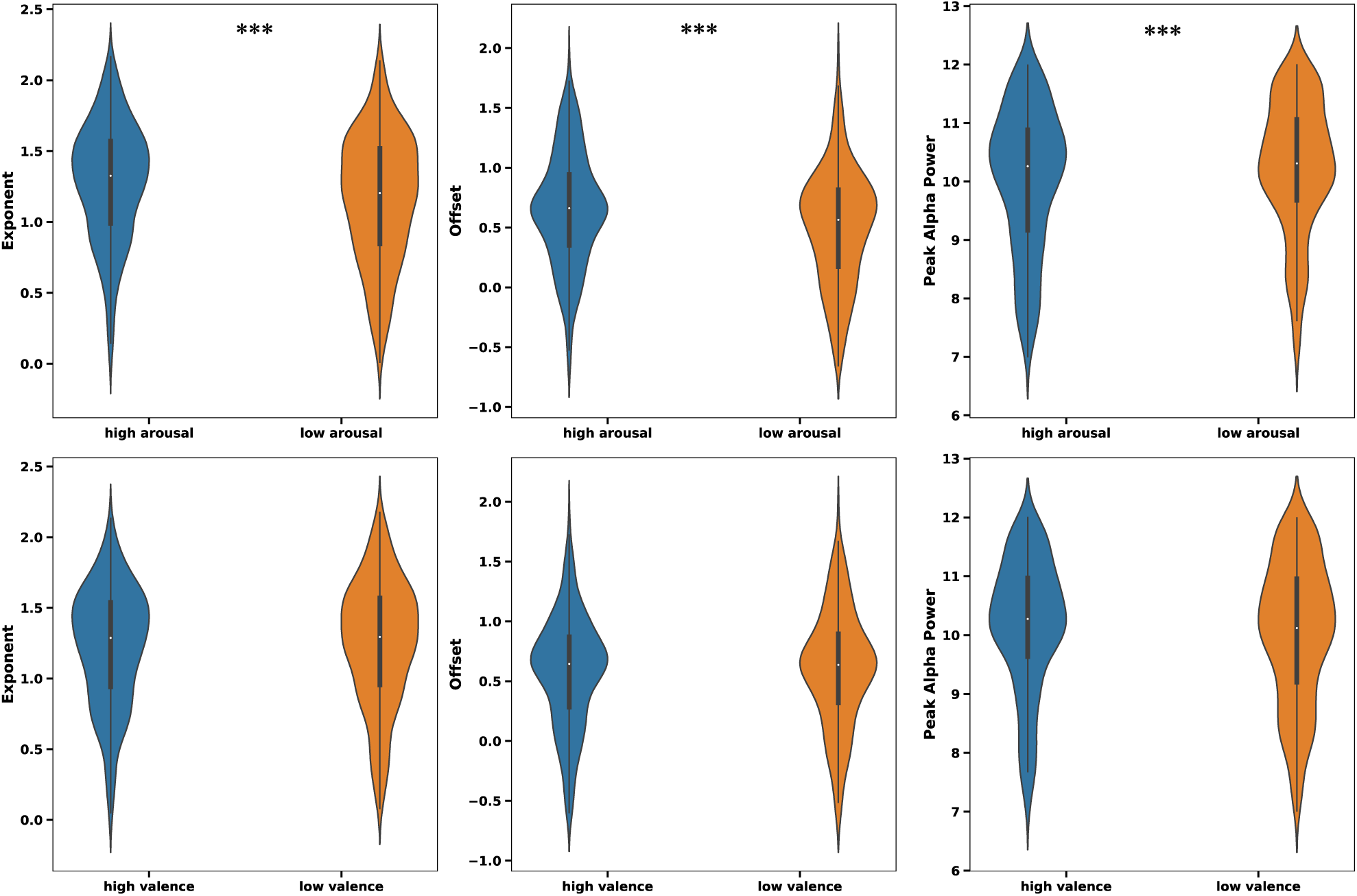
Aperiodic spectra and components between conditions [***p<0.001, **p<0.01, *p<0.05]. Power in μV^2^, Frequency in Hz, Exponent in μV^2^Hz^-1^

**Table 2:**
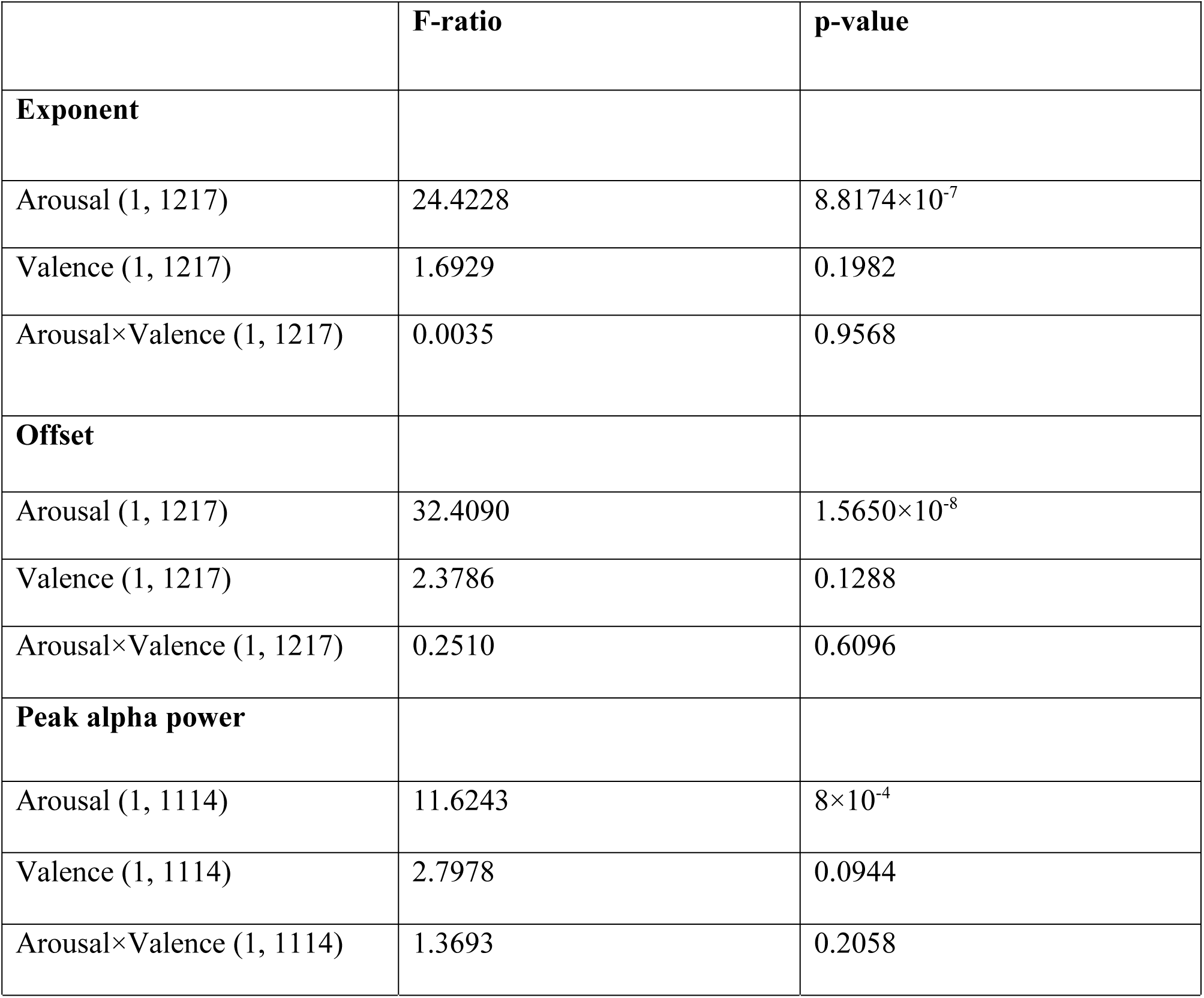
Two way random permutation ANOVA results on exponents and offsets of 1/f slope and peak alpha power with arousal and valence as the factors

|  | **F-ratio** | **p-value** |
| --- | --- | --- |
| **Exponent** |  |  |
| Arousal (1, 1217) | 24.4228 | $8.8174 \times 10^{-7}$ |
| Valence (1, 1217) | 1.6929 | 0.1982 |
| Arousal×Valence (1, 1217) | 0.0035 | 0.9568 |
| **Offset** |  |  |
| Arousal (1, 1217) | 32.4090 | $1.5650 \times 10^{-8}$ |
| Valence (1, 1217) | 2.3786 | 0.1288 |
| Arousal×Valence (1, 1217) | 0.2510 | 0.6096 |
| **Peak alpha power** |  |  |
| Arousal (1, 1114) | 11.6243 | $8 \times 10^{-4}$ |
| Valence (1, 1114) | 2.7978 | 0.0944 |
| Arousal×Valence (1, 1114) | 1.3693 | 0.2058 |

A significant increase in the average exponent value (*Δ Χ* _(ℎ_*_ig_*_ℎ_*_arousal − lowarousal_*_)_ = 0.1197 ± 0.0123, p = 3.2780×10^-4^, effect size = 0.2519) and a significant increase in offset (*Δb*_(ℎ_*_ig_*_ℎ_*_arousal− lowarousal_*_)_ = 0.1582 ± 0.0144, p = 1.9996×10^-4^, effect size = 0.3083) were observed as arousal levels increase. Furthermore, we noted a significant decrease in the average peak alpha power with arousal (*Δa*_(ℎ_*_ig_*_ℎ_*_arousal − lowarousal_*_)_ = -0.2400 ± 0.0342, p = 0.0097, effect size = -0.1767).

On the other hand, the exponent value exhibited no significant changes across trials reported as low and high valence (*Δ Χ* _(ℎ_*_ig_*_ℎ_*_valence −lowvalence_* _)_ = -0.0057 ± 0.0099, p = 0.9203, effect size = -0.0098), and similar trends followed for offset (*Δb*_(ℎ_*_ig_*_ℎ_*_valence −lowvalence_* _)_ = -0.0119 ± 0.0139, p = 0.8630, effect size = -0.0043) and average alpha peak power (*Δa*_(ℎ_*_ig_*_ℎ_*_valence −lowvalence_* _)_ = 0.0687± 0.0301, p = 0.6086, effect size = 0.0328).

### Spectral modulations with stimulus presentation in a thalamocortical model of EEG

A thalamocortical model was set up to probe the mechanisms underlying changes in periodic and aperiodic properties of ongoing EEG activity with emotional stimuli. In the absence of stimuli, the relay populations are driven by noisy inputs, and the resulting power spectra correspond to a resting awake state (see Figure 5). From a dynamical systems point of view, the activity of the cortical population is akin to a damped oscillator with a natural frequency lying in the alpha range (8-12Hz). The input to the relay cells is modeled as a random input derived from a Gaussian distribution (mean =0, std = 1).

**Figure 5:**
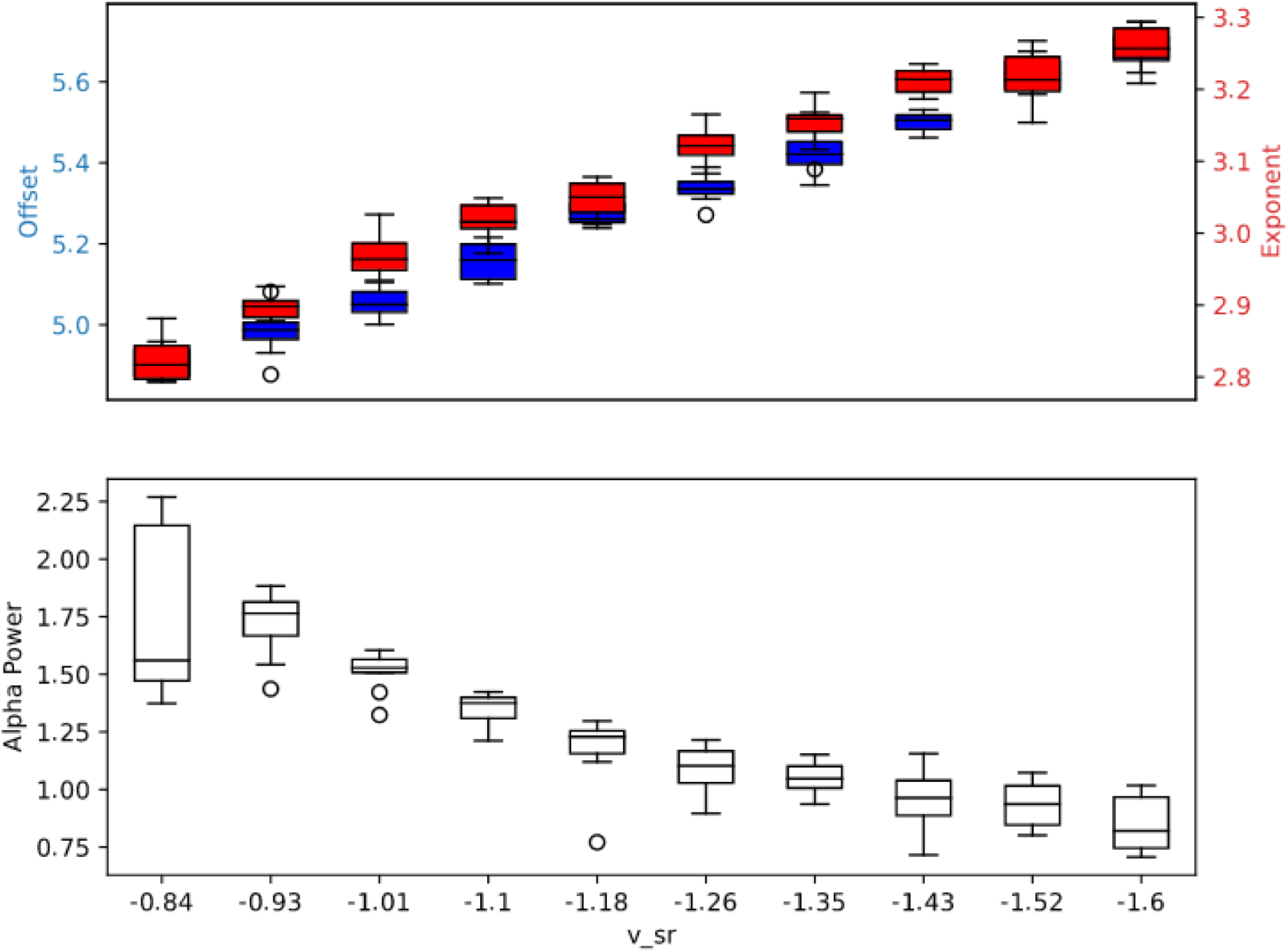
Aperiodic components and Aperiodic power with change in inhibitory strength from reticular to relay nucleus of thalamus (v_sr)

The arousal condition is simulated by increasing the inhibitory coupling strength from reticular to relay thalamic population. We perform numerical integration of equation (4) with Euler method and step size of 1ms. Transients were removed from the stimulated time series, following which the resulting spectra is modeled with the FOOOF algorithm (same parameters as the EEG analysis) to extract periodic and aperiodic features, namely the peak alpha power and the offset and exponent of the background activity. We could observe that stimulus presentation results in a reduction of alpha power, a steepening of the 1/f slope and an increase in the offset, similar to what is seen empirically. Further, the magnitude of the effect scales monotonically with the stimulus intensity (see Figure 5).

## Discussion

The primary aim of our study was to investigate how emotional states conceptualized along dimensions of arousal and valence is represented in EEG signals in a naturalistic setting. In this context we found that the contributions of periodic and aperiodic background is scantly researched in the context of dimensions of emotion. With the current understanding in the field that rhythmic and arrhythmic activity can be stemming from distinct neural processes (Donoghue et al., 2020; Voytek et al., 2015), we hypothesized that arousal and valence may be differentially represented along these signal attributes. Our findings demonstrated a novel significant increase in both parameters that define the aperiodic component, namely, exponent and offset during the viewing of video clips perceived as eliciting high arousal, in contrast to those with low arousal. This result suggests a potential connection between the aperiodic components and the intensity of emotional arousal, expanding our understanding beyond the commonly studied rhythmic components. Surprisingly, we did not observe significant differences in aperiodic components when comparing high and low valence classifications, suggesting that the influence of valence on these EEG features may be subtler. The decrease in peak alpha power during high arousal compared to low arousal conditions aligns with previous studies linking alpha power to emotional arousal (Hofmann et al., 2021; Kim et al., 2021; Schubring & Schupp, 2021). Digging deeper, to uncover the mechanistic underpinnings of the arrhythmic and rhythmic constituents, our investigation utilized the thalamo-cortical mode (Robinson et al., 1997, 2003). Notably, our results suggest that increased coupling between thalamic reticular and relay nuclei within this model can replicate the pattern of change in aperiodic and periodic activity with increasing arousal. The coupling is of inhibitory nature, hence it supports the inhibitory gating of stimuli from amotion arousal which we expand in the follow up section.

### Increased inhibition with emotional arousal

The exponent of 1/f component, exhibits an increase during high arousal states compared to low arousal states. This exponent is the negative counterpart of the slope within the equations utilized by our algorithm. In the context of aperiodic spectra, the slope has been proposed to represent a balance between excitation and inhibition, as demonstrated in previous studies (Gao et al., 2017) (Miller et al., 2009) (Voytek et al., 2015) Thus, a steeper slope (more negative) or a larger exponent suggests a greater degree of inhibition relative to excitation. Our results consistently indicate a higher exponent during high arousal, signifying enhanced inhibitory processes during heightened emotional states. This observation suggests that high emotional arousal may involve active inhibitory control, marking a departure from certain studies that have linked arousal, defined in terms of vigilance levels, e.g. command following, eyes opening; (Boly et al., 2013) during conscious states of sleep, anesthesia, and wakefulness to the 1/f slope (Colombo et al., 2019; Lendner et al., 2020). Notably, the slope was found to be more negative during sleep stages compared to the awake state. It is important to emphasize that different arousal types are defined in Neuroscience literature; sleep-wake, alertness level, physiological, sexual, emotional (e.g. (Pfaff et al., 2012)(Hofmann et al., 2021; Pfaff et al., 2012) (Hofmann et al., 2021). Arousal in the context of sleep studies typically pertains to the non-specific activation of the cortex within the sleep-wake dimension (Oken et al., 2006). Nevertheless, they are different processes and yet may not be completely dissociable; they share some psychological (e.g. enhanced sensorimotor activation, emotional reactivity) and physiological (e.g. sympathetic activations) characteristics (Kreibig, 2010) (Pfaff et al., 2012) (Hofmann et al., 2021). On this account, the reversal of the exponent value observed in our study suggests that the mechanism of arousal in emotional contexts may differ from arousal within the sleep-wake dimension.

### Increased offset and reduced alpha power with increased arousal

The aperiodic offset, can give a qualitative description of average firing rate of neuronal populations (Manning et al., 2009; Miller et al., 2009) (Miller et al., 2009) and can be conceptualized as an indicator of the overall neural activity or cortical excitability(Weber et al., 2019). Weber et al., 2019, reported a decrease in broadband power (defined by the offset estimation) during isolation,(6 member crew underwent a 120-day isolation period, simulating conditions analogous to space travel) suggesting a reduction in overall neural activity. Consequently, this interpretation implies an elevated level of cortical activation during high arousal in comparison to low arousal settings. Additionally, the reduction in alpha peak power, which aligns with prior research, is observed during high arousal states. This decline in alpha power has been proposed to result from heightened attentional demands (Ghosh et al., 2021; Hofmann et al., 2021; Jensen & Mazaheri, 2010; Kim et al., 2021; Schubring & Schupp, 2021; Weisz et al., 2011; Wöstmann et al., 2021) .

### A plausible mechanism of emotion attention modulation at thalamo-cortical level

In light of the primate studies demonstrating direct connections between the amygdala and the thalamic reticular nucleus, particularly in the context of emotion and selective attention (John et al., 2016; Zikopoulos & Barbas, 2012), and the role of the thalamic reticular nucleus in sensory processing, as shown in studies involving rats (Ahrens et al., 2015; Aizenberg et al., 2019; Shosaku, 1986), we sought to translate these findings along with our observations into a computational model of the thalamocortical system. The model was designed to receive noisy ascending inputs and, intriguingly, behaves akin to a damped oscillator, returning to baseline activity following perturbations at a characteristic frequency. In our model, we use the mean membrane voltage of excitatory cortical populations to represent cortical activity. Importantly, the resting dynamics of our model encompass key variables of interest in our study, including a prominent spectral peak in the alpha band and distinctive 1/f background activity, as previously observed in empirical studies (Robinson et al., 1997, 2003). Modulating the inhibitory coupling strength from the reticular thalamocortical unit allowed us to explore how the spectral statistics of the alpha band and background aperiodic activity, aligns with empirical observations. This we hypothesize, could be a result of amygdala to thalamic reticular connection. However, in humans, evidence of direct connections has been reported between amygdala and pulvinar thalamic nuclei, (Abivardi & Bach, 2017; Tamietto et al., 2012) and also been suggested as a pathway for emotion attention (Pessoa & Adolphs, 2010; Vuilleumier, 2005). With respect to this, another possible modulation can be from amygdala to relay nucleus will have to be inhibitory in nature. This requires however, a detailed investigation in future studies with direct anatomical measurements currently out of scope of the present data set.

### Change in aperiodic components with a sensory input

Furthermore, we noticed an increase in both the aperiodic offset and exponent during video presentation, irrespective of valence and arousal categorization (Supplementary Figure, S2). This observation suggests a dynamic interplay between aperiodic activity and sensory input during the presentation of visual stimuli. Interestingly, (Weber et al., 2019) reported that following the discontinuation of the isolation period, broadband power returns to its pre-isolation levels. This underscores the influence of sensory stimulation and deprivation on broadband power dynamics, Notably, (Gyurkovics et al., 2022) found that following an auditory stimulation, an increase in 1/f activity was observed compared to the pre stimulus state. Moreover, the steepness of the slope was increased with rare oddball stimuli compared to standard auditory stimuli. They thus proposed correlation of aperiodic slope with the attentional orientation through inhibitory process. Complementing the aperiodic slope relation to attention shift and cognitive processing, (Ostlund et al., 2021) reported flatter slope in ADHD (inefficient information processing) adolescent population compared to age matched non-ADHD population. Thus, these studies provides an interesting hypothesis of aperiodic slope being as a marker for attentional shift and possibly an information processing tendency. Therefore, the increased offset and exponent during stimulus presentation compared to pre stimulus state could reflect attentional demand of the emotional stimuli here.

### Limitations and future directions

In examining emotional responses to videos, various factors impact the analysis methodology and its interpretation. Firstly, the study is constrained by assessing subjective emotions only after the video concludes. Thus we assume that reported and felt emotional states remain constant for throughout that one minute of video presentation. However, it’s important to acknowledge that emotions may evolve during this period, and the fixed assessment may not fully capture potential changes. Consequently, this doesn’t allow for temporal modelling of thalamic gating of emotion. Additionally, the model assumes the cortex as an interaction of a single excitatory and inhibitory population, lacking consideration for spatial specificity. This limits understanding of the spatio-temporal dynamics of emotional experiences during video exposure. To comprehensively evaluate the results, it is crucial to interpret them within the context of continuous rating and on a subject-specific basis. Thus, it is essential to recognize that individuals experience emotions uniquely and that emotional responses can fluctuate over time.

