## Supplement figure for "Emotion arousal but not valence is strongly represented in aperiodic EEG activity stemming from thalamocortical interactions"

### Supplementary Results

#### Spatial topography of exponent, offset and residual alpha

To assess the spatial distribution of the spectral components, we plot the difference in exponent, offset and residual power (obtained by subtracting the aperiodic fit from spectrum model fit). We find exponent is higher in frontal electrodes implying more inhibition of the frontal areas compared to low arousal. We also find increased offset represented at the frontal electrodes with arousal, indicating increased neural activity. Thus, there is an increase in frontal neural activity but as an inhibitory modulation. And as observed in previous studies (e.g. (Kim et al., 2021; Lang et al., 1998)), we find decreased alpha power in posterior regions. This has been inferred to be heightened sensory processing to increase attention demand.

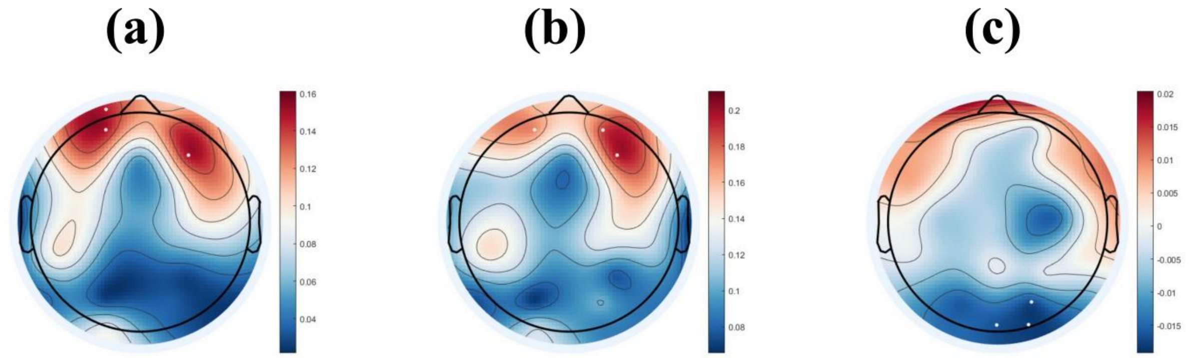


**S1: Differences between high arousal and low arousal trials are represented and electrodes showing >0.9 differences are shown in white (a) is difference between exponents (b) is difference in offset (c) is difference in residual alpha**

#### Increase in exponent and offset with incoming stimulus independent of arousal and valence categorization (60s video signal length)

We performed paired two tailed sign test (signtest.m) was to show change in aperiodic components in task from pre stimulus condition. The test takes difference of the two samples in consideration and finds if values are greater or less than zero or is equal to zero. The values are then assigned signs of +1, -1, 0 based on the positive, negative and no difference respectively.

We found there is a significant increase in exponent in task state of high arousal trials compared to pre stimulus of high arousal trials ($Z = 7.9904$, $p < 0.0001$, positive $sign_{(task-pre)}$=470 out of 724); similar results were also observed on comparing low arousal pre stimulus and task state (low arousal, $Z = 3.4845$, $p < 0.0001$, positive $sign_{(task-pre)}$=297 out of 514). We also repeated the procedure for valence trials and found the same observation (high valence, $Z = 5.5561$, $p < 0.0001$, positive $sign_{(task-pre)}$= 424 out of 700; low valence, $Z = 6.0677$, $p < 0.0001$, positive $sign_{(task-pre)}$=341 out of 540).

In addition we also compared offsets of pre and task state in the similar manner and found an increase in offset value that from the pre stimulus state in both categories (high arousal, $Z = 11.7069$, $p < 0.0001$, positive $sign_{(task-pre)}$= 520 out of 724; low arousal, $Z = 9.5715$,$p < 0.0001$, positive $sign_{(task-pre)}$= 366 out of 514; high valence, $Z = 12.6618$, $p < 0.0001$, positive $sign_{(task-pre)}$= 518 out of 700; low valence, $Z = 8.9074$, $p < 0.0001$, positive $sign_{(task-pre)}$= 374 out of 540).

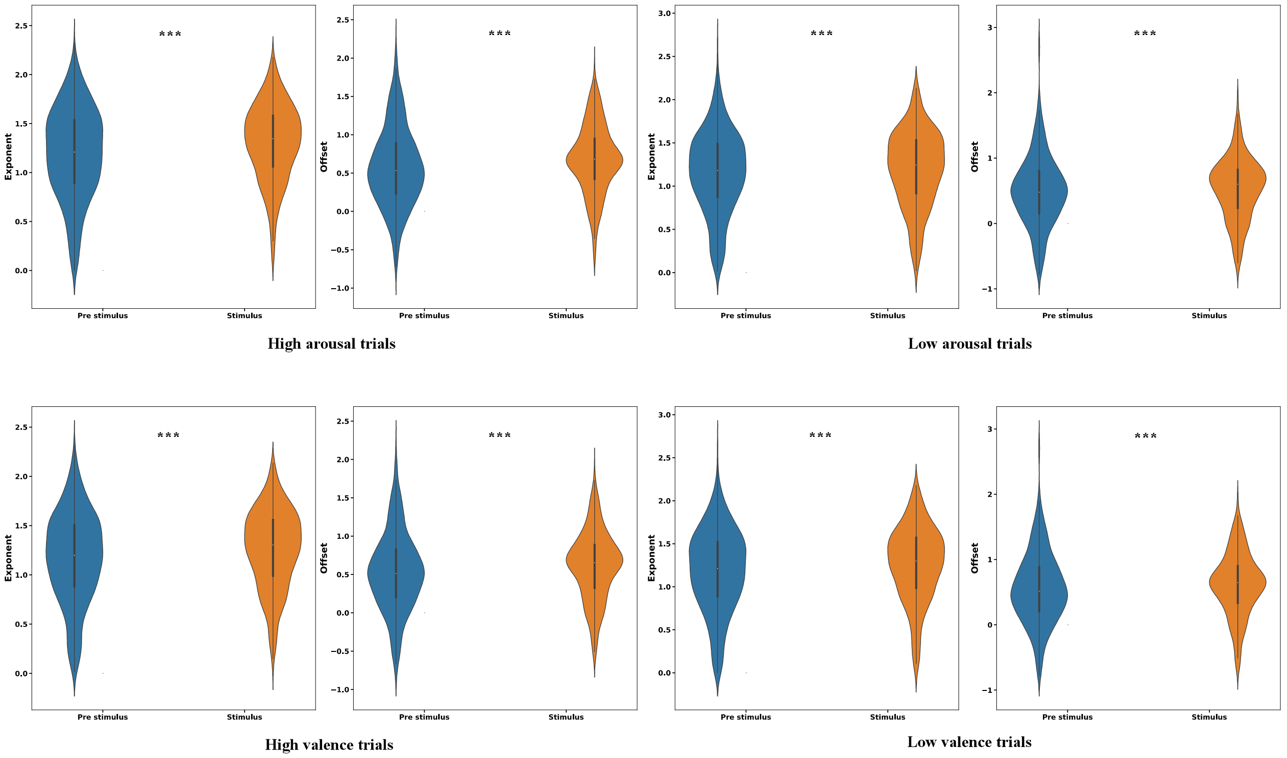


**S2: Pre stimulus and stimulus aperiodic components in arousal and valence trials(***p<0.001, **p<0.01,*p<0.05)**
